# Comparison of a Novel Real-World Speech in Noise Auditory Attention Task to Standard Clinical Audiological Metrics

**DOI:** 10.64898/2026.05.28.727658

**Authors:** Brett M. Bormann, Nina E. Wade, Kelsey Mankel, Daniel C. Comstock, Soukhin Das, Richard S. Whittle, Hilary Brodie, Doron Sagiv, Lee M. Miller

## Abstract

Pure tone audiometry (PTA) remains the clinical standard for evaluating hearing ability, yet individuals with similar audiometric profiles often exhibit substantial variability in their capacity to understand speech in everyday listening environments. Growing evidence suggests this variance is related to contributions from cognitive ability and auditory processing that standard threshold measures do not capture. To investigate how PTA, cognitive factors, and demographics such as age jointly predict real-world speech perception, 116 veteran adults 20-70 years old spanning a range of normal to moderate sensorineural hearing losses completed a spatial auditory attention task. Target color words were embedded within naturalistic short-story narratives presented under two conditions: a mono-talker speech-in-quiet (SIQ) condition and a dual-talker speech-in-noise (SIN) condition with a spatially separated competing narrative. Behavioral performance was quantified via color word hit accuracy, reaction time, and comprehension question accuracy. Participants also completed pure tone audiometry, the Montreal Cognitive Assessment (MoCA), and the Speech, Spatial and Qualities of Hearing Scale (SSQ12). Mixed-effects regression models were used to evaluate the contributions of PTA, age, cognitive ability, and self-reported hearing difficulty (SSQ12) to task performance across conditions. Results demonstrate a complex interplay between age, PTA, MoCA, and/or listening condition (SIQ vs. SIN) in predicting identification accuracy, reaction time, and comprehension. Age and condition significantly predicted hit accuracy and reaction time, with older participants showing improved accuracy in quiet but declining accuracy and slower responses in noise. PTA did not emerge as a significant main effect predictor but interacted with cognitive ability and condition to modulate performance, in some cases exhibiting a paradoxical inverse relationship with accuracy dependent on MoCA score. MoCA scores significantly predicted comprehension across conditions, and SIN hit accuracy was positively correlated with SSQ12 scores, validating the task against participants’ real-world listening experiences. These findings highlight the importance of incorporating cognitive screening and ecologically valid speech perception tasks into audiological assessment to better identify individuals at risk for functional hearing impairment in complex listening environments.

## Introduction

Hearing loss is a significant public health concern, affecting over 16% of adults in the United States and approximately 432 million people worldwide with disabling hearing loss (>40 dB HL) (Carroll, 2017; *WHO: Deafness and Hearing Loss*, n.d.). Hearing ability is critical for social, psychological, and cognitive aspects of life (Henderson et al., 2025), and undiagnosed or under-assessed hearing difficulties can have serious implications for quality of life, cognitive decline, and comorbidities (*Audiology Information Series: Comorbidities and Hearing Loss*, 2021; Besser et al., 2018). Notably, 10-15% of patients presenting with clinically “normal” audiograms report difficulties hearing in noise, a phenomenon referred to as hidden hearing loss because their perceptual challenges are “hidden” from traditional auditory assessments (Parthasarathy et al., 2020; Spankovich et al., 2018; Tremblay et al., 2015). These observations highlight a growing concern that commonly used clinical measures may fail to capture meaningful variance in real-world listening ability, particularly under acoustically challenging conditions (Cord et al., 2007; Dubno, 2018; Nuber, 2016). Accurate, sensitive, and comprehensive diagnostic criteria are therefore essential for identifying functional hearing deficits that may not be captured by standard clinical metrics.

Pure tone threshold audiometry remains the gold standard for evaluating hearing in clinical settings (Salmon et al., 2023; Tanna et al., 2025). Pure tone thresholds quantify the quietest sound an individual can detect 50% of the time across a range of frequencies, typically 250-8000 Hz, to determine the type and degree of hearing loss (American Speech-Language-Hearing Association, n.d.; Impairments et al., 2004; Walker et al., 2013). Thresholds below 25 dB HL are considered clinically normal (American Speech-Language-Hearing Association, n.d.; Salmon et al., 2023), with some groups starting to recognize a “slight” hearing loss category for thresholds between 16-25 dB HL (American Speech-Language-Hearing Association, n.d.). Disabling hearing loss is generally defined as thresholds exceeding 35 dB HL (*WHO: Deafness and Hearing Loss*, n.d.). Pure tone averages (PTAs), typically calculated as the average of thresholds at 500, 1000, and 2000 Hz, summarize hearing within the speech-relevant frequency range (Salmon et al., 2023). Audiograms and PTAs are heavily relied upon due to their objectivity, efficiency, and ability to extract patterns of auditory sensitivity (Salmon et al., 2023). However, while pure-tone audiometry remains a foundational clinical tool, it has notable limitations in assessing the complex auditory processes required for real-world speech perception, particularly in acoustically challenging environments (Musiek et al., 2017). As a result, PTA may capture only a limited portion of the variance underlying functional communication ability in everyday listening environments.

Real-world speech perception requires not only intact auditory sensitivity but also efficient integration of auditory, neural, and cognitive elements. Competing inputs, overlapping speech, and environmental noise demand that listeners isolate relevant sounds, encode them with temporal and spatial precision, and integrate linguistic information in context (Alain & Arnott, 2000; Giovanelli et al., 2024; Pichora-Fuller et al., 1995). Furthermore, difficulties in speech comprehension in noise are observed even in the absence of measurable hearing impairment. Many individuals report difficulty understanding speech-in-noise even when audiometric thresholds fall within normal ranges (Alain C & Tremblay K, 2007; Bidelman et al., 2019; B. A. Schneider et al., 2002; Tremblay et al., 2015), suggesting that factors beyond audibility contribute substantially to performance in complex listening environments. This may stem from cognitive and neural influences that standard audiometry cannot detect (Barbee et al., 2018). This underscores the need for assessment approaches that extend beyond quiet, threshold-based measures to better characterize real-world hearing difficulties. Thus, while pure-tone audiometry provides valuable standard measures for clinical assessment, additional factors contributing to complex auditory perception are not captured by this test alone (S. Anderson et al., 2013). While clinical speech and speech-in-noise tests exist as supplements to pure tone audiometry, their use in routine practice remains inconsistent, and when administered, the stimuli and conditions may fall short of capturing the acoustic complexity of everyday listening environments (M. C. Anderson et al., 2018; Billings et al., 2023; Parmar et al., 2022).

Aging adults face additional challenges, as age-related auditory and cognitive changes interact to impair speech perception (S. Anderson et al., 2013). Mechanical and neural degradation along the auditory pathway, from the outer ear to the auditory cortex, can reduce processing efficiency even before hearing loss is clinically measurable (Anastasiadou & Al Khalili, 2025; Morrell et al., 1996; B. Schneider, 1997). These sensory declines are often accompanied by age-related declines in cognitive domains critical for speech comprehension (Wingfield & Stine-Morrow, 2000). Such age-related effects do not manifest uniformly across individuals, and perceptual and cognitive factors interact dynamically over the lifespan, producing variable and sometimes subtle changes in speech perception that may not be reflected in PTA alone.

The goal of this study is to evaluate the sensitivity and reliability of PTA in predicting speech-in-quiet (SIQ) and speech-in-noise (SIN) perception in a cohort of veteran adults spanning a wide range of ages and audiometric profiles. Participants completed a naturalistic, continuous speech task under two conditions—mono-talker (speech-in-quiet; SIQ) and dual-talker (SIN)—where they identified embedded color words as a behavioral measure of listening performance.

Participants were also evaluated on their comprehension of the short story narratives. We hypothesized that PTA would explain only a small proportion of variance in SIN performance, even after accounting for age, and that its predictive value would be weaker for SIN than SIQ. Our findings aim to clarify the extent to which traditional audiometric thresholds reflect real-world listening ability and whether current diagnostic practices are sufficient for identifying individuals at risk for functional hearing impairment in complex auditory environments.

## Materials and Methods

### Participant Information

We collected data from 116 veteran participants aged 20-70 years old (mean age = 45.52 years, SD = 13.91; 84 males, 32 females). Participants were veterans who were off duty, on leave, or retired at the time of the study. All were fluent in English and had normal or corrected to normal vision (<20/40 on Snellen chart). None had been diagnosed with severe or profound hearing loss or used assistive listening devices such as hearing aids or cochlear implants. Audiometric evaluation confirmed that participants had varied hearing ranging from normal hearing to moderate sensorineural hearing loss. Inclusion criteria required pure tone air conduction thresholds of <60 dB HL in the better ear across the 250-4000 Hz range, an air-bone gap of ≤10, and ≤20 dB interaural asymmetry at 500-2000 Hz. All participants completed a cognitive screening to ensure no cognitive deficits that could act as a confound using the Montreal Cognitive Assessment (MoCA), with a score of 24 or higher (Nasreddine et al., 2005). None reported a history of brain injury or neurological/psychiatric conditions that would directly affect their ability to pay attention or understand speech (e.g., epilepsy, stroke, or ADHD). All participants completed an informed consent according to a protocol approved by the University of California, Davis Institutional Review Board and were financially compensated for their participation and transportation.

### Project Overview

Participants completed three sessions over one, two, or three days, with breaks throughout to reduce fatigue. The sessions included a one-hour audiological session, a one-and-a-half-hour cognitive-behavioral session, and a two-and-a-half-hour electroencephalography (EEG) session. The following analysis uses a portion of the data collected from this larger study, specifically focused on clinical audiological measures and performance on the spatial speech perception task from the EEG session.

## Audiologic Session

Participants completed a standard audiologic assessment either at the UC Davis Otolaryngology and Audiology Clinic administered by a professional audiologist or in a sound-dampened room at the Speech Neuroengineering and Cybernetics Lab at UC Davis by a trained lab researcher.

### Otoscopy

Audiologists or lab researchers performed otoscopy to visualize the outer ear and tympanic membrane, screening for abnormalities, occlusions, or any pathology that would prevent the use of inserts. If any pathologies or occlusions were noted, participants were encouraged to resolve the issue with professional care before returning to the study.

### Pure Tone Audiometry

Pure tone audiometry was conducted for both ears (GSA Audiostar Pro or Inventis Cello). Air conduction thresholds were obtained using pulsed tones delivered through insert earphones at 250, 500, 1000, 2000, 3000, 4000, 6000 and 8000 Hz (Radioear IP30 or ER-3C). Pure tone average thresholds (PTA) were calculated as the mean threshold across 500, 1000, and 2000 Hz for each ear. Extended high frequency (EHF) thresholds were also collected at 9000, 10000, 11000, 12000, 16000, and 18000 Hz using specialized Radioear DD450 headphones. Bone conduction thresholds were collected at 250, 500, 1000, 2000, 3000 and 4000 using pulsed tones via a bone conductor placed on the mastoid of the ear with the lower PTA (Radioear B81 or B71). All thresholds were obtained using a standardized 10 dB down / 5 dB up staircase procedure.

### The Speech, Spatial and Qualities of Hearing Scale

Participants completed the Speech, Spatial, and Qualities of Hearing Scale (SSQ12), a standardized 12-item self-assessment questionnaire designed to capture hearing-related functional abilities across three domains: speech perception in complex listening environments, spatial hearing (e.g., direction, distance, movement), and perceptual qualities of hearing such as listening effort, clarity, and naturalness (Gatehouse & Noble, 2004). The SSQ12 asks participants to rate real-world listening scenarios using a Likert-type scale, with higher scores reflecting better perceived performance. The SSQ12 provides insight into how individuals perceive their hearing ability in everyday listening contexts, capturing functional deficits that may not be evident from audiometric thresholds alone. For the purposes of this study, we used the overall SSQ12 score, calculated as the mean of all item ratings across the three subscales.

### Additional Audiometric and Behavioral Assessments

Although not included in the present analyses, we also collected audiological data including tympanometry, acoustic reflexes, distortion product otoacoustic emissions (DPOAEs), speech recognition thresholds, and word recognition scores. However, these measures are out of scope for the current analyses which specifically focus on pure tone audiometry. Additional cognitive tests were collected during the behavioral session: SPRINT, Stroop, Reading Span, Temporal Fine Structure, Flanker test, TMT, pitch discrimination. These were also out of scope for the current analyses. However, recent literature increasingly supports the utility and correlation between cognitive faculties and auditory perception (Shehabi et al., 2025).

## Continuous Multi-Talker Spatial Auditory Attention Task

### Stimuli

Details of the stimuli are specified in additional reports from this study (Mankel et al., 2025; Shehabi et al., 2025) but briefly reviewed here. Participants listened to six 7.5-minute passages of short-story narratives drawn from public-domain fairy tales with content unlikely to be recognized as modern renditions. Stories were selected based on engaging content, similar story arc progress, ease of understanding, and grammar complexity for an adult audience as rated by four pilot listeners prior to the study. Each story was modified to naturally embed monosyllabic target color words within the story (e.g., blue, black, white, etc.). The stories were recorded in a soundproof booth using a Shure KSM244 vocal microphone (cardiod polar pattern, high-pass filter at 80 Hz with 18 dB per octave roll-off) and Adobe Audition at a 48 kHz sampling rate. All were narrated by a male, native English speaker with an American Midwestern accent. Silent gaps in the recordings greater than 500 ms were shortened to 500 ms, similar to Broderick et al. (2018), Teoh et al. (2022), and Teoh & Lalor (2019), and the trimmed recordings were then cropped to 7.5 minutes per story audio.

The audio stimuli were additionally modified with “Cheech,” a patented chirped-speech process in which glottal pulse energy is replaced with narrowband synthetic chirps (Backer et al., 2019; Miller & Moore, 2020). Cheech preserves naturalistic speech and linguistic content while enabling simultaneous measurement of auditory evoked potentials across the brainstem, thalamus, and cortex, offering greater control over the acoustic parameters of speech used to evoke these responses. Although Cheech’s neurolinguistic applications are not used here, relevant references describe its broader utility (Backer et al., 2019; Corina et al., 2022, 2024; Miller & Moore, 2020; Shehabi et al., 2025).

Of the six passages, two simulate female talkers by resynthesizing the original male voice, shifting the mean F0 from 128 Hz to 180 Hz and using a bilinear transform to warp vocal tract length (Zhan & Waibel, 1997), with warping factor (alpha) of 1.04 up to a maximum frequency of 4000 Hz. Half of the stories were modified with standard Cheech parameters, and three stories were subjected to an additional experimental manipulation designed to affect neural encoding (which are not used in this study. One additional demo story was used for participant orientation to the stimuli and practice of the task. Of the six story blocks that the participants completed, this analysis focuses on the following male talker target conditions: a mono-talker condition, where a single story was presented, and a dual-talker condition, where a target and masker story were presented simultaneously. Each of the two conditions were comprised of a single-story block, so two of the six blocks are used for this analysis, excluding dual talker female target stories for parity (i.e., avoid confounds associated with male vs. female voice (Mankel et al., 2025)). These conditions were selected specifically to focus on the impacts of an ecologically relevant masking noise on the participants’ speech perception.

### Experimental Conditions

Auditory stimuli were spatially filtered using head-related transfer functions (HRTFs) from the SADIE II database (Armstrong et al., 2018) to stimulate audio at ±15°, which was mirrored on two computer screens centered around 15° left and right of the participant’s midline. Participants sat in a soundproof booth approximately 176 cm away from the computer screens. In the dual-talker paradigm, each story was presented twice, once as the target story and once as the masker story within its pair. This design allows us to assess the effects of directed spatial attention, as participants were instructed to either attend to or ignore the same audio depending on its role in each trial. In dual-talker blocks, the target story was presented from one spatial location and the masker story from the opposite side, whereas in the single-talker blocks, the target story was delivered from only one side at a time. In both cases, the left/right assignments switched multiple times during the block at each side switch, both the spatialized audio and the on-screen cue transitioned to the new target location, and participants were instructed to immediately shift their gaze and auditory attention accordingly. The cue on screen indicated location of the target story as being in the left or right spatial location with a “<” or “>” symbol, and a “+” symbol cueing the target location. Auditory and visual cues switched sides approximately every six seconds (75 total switches), with visual symbols matched in size, color, and line length. Stimuli were presented using custom MATLAB and Psychtoolbox scripts. Audio was routed through a Hammerfall audio card, RME Fireface UFX II interface, and RME ADI-2DAC FS amplifier, and delivered via ER-2 insert earphones with shielded wires to minimize EEG artifacts. Cheech stimuli were presented at 75 dB SPL. Color word onsets occurred between 0 and 2 seconds following a spatial switch. Visual icons updated instantaneously at the switch, and the audio transitioned with a 35ms crossover fade starting at the switch cue. Each block contained 25 color words, which were randomly assigned to one of the five following onset times relative to the switch (0.125, 0.25, 0.5, 1 or 2 seconds; 5 words per condition).

### Procedure

The procedure was monitored and presented by custom MATLAB and Psychtoolbox code. Participants first completed a demo block to familiarize themselves with the task and Cheech-processed audio. There was a 1-minute resting state period at the start of each block. Each story block included 7.5 minutes of story in which participants were instructed to listen to the story, attending to the target voice and ignoring the distractor story when applicable. They were told to press the spacebar with their right hand when they heard a color word in the target story while ignoring any target words presented in the distractor story (if present). After each story, participants were prompted to answer 5 multiple-choice comprehension questions, each with four answer choices. Pilot testing calibrated questions ensuring a balance in difficulty (neither too hard nor easy) and assessed both memorization and broader inferences. Lastly, participants completed a visual reaction time task by pressing the spacebar in response to a white circle flash, used as a measure of fatigue. Story presentation order was randomized using a Latin square design.

### Behavioral Measures

Accuracy in detecting target color words served as a behavioral indicator of speech-in-quiet (SIQ) perception and speech-in-noise (SIN) perception. A color word hit constituted pressing the spacebar within 2 seconds after the onset of the color word in the target story. Target words that occurred within ±2 seconds of a color word in the masker story were excluded, yielding 21 target color words for the SIN condition and 25 for the SIQ condition. Accuracy was calculated as the number of successful hits divided by the number of eligible target color words per block.

Reaction time was also captured for each color word hit. Following each block, participants answered comprehension questions in which average accuracy across the five questions was calculated. Neural measures from the EEG are not analyzed here but instead included in other publications from this study (Mankel et al., 2025; Shehabi et al., 2025).

## Statistical Analysis

All statistical tests were performed with R version 4.5.2 (Posit Software, PBC) on a computer running macOS Tahoe 26.2. Any observations that were more than 3 standard deviations from the mean of all the observations of a given variable were labeled as outliers. A Cook’s distance analysis was performed for each of those observations to determine if they were influential by the calculated cook distance being greater than 4 divided by the number of observations. Any observations that were identified as influential outliers were removed. Any missing values due to outlier removal or technical issues during data collection were imputed using non-linear interactive partial least squares (“Nonlinear Iterative Partial Least Squares (NIPALS) Modelling,” 1973).

Pairwise t-tests were used to check for significant difference between task performance (color word hit accuracy, color word hit reaction time, and comprehension question accuracy) across the SIQ and SIN conditions.

Mixed modeling multiple regression was used to evaluate the relationship between participant’s performance on our novel spatial attention auditory task with multiple variables that we believe relevant for evaluating a person’s listening ability. A model was created for each performance metric (color word hit accuracy, color word hit reaction time, and comprehension question accuracy) where their performance was predicted by the six variables (task condition (SIQ or SIN), participant’s age, PTA values, SSQ12 scores, MoCA scores, and NASA task load values). All predictor variables were scaled using Z-score normalization (but plotted using raw data values). These initial models included every variable and interaction between the variables. The models were then optimized using recursive feature elimination to remove any nonsignificant interactions. Mixed effects beta multiple regression was used to model the participants’ color word hit and comprehension question accuracies because those values were bound from 0-100%. The participants’ reaction time were not bound so mixed effects linear multiple regression was used. Significance testing of these models was performed with a type 3 ANOVA. Marginal means were then used to test for the significance of individual predicted model trends.

## Results

### Audiometric Measures

116 participants (mean age = 45.52, SD = 13.91) completed a pure tone audiogram where their pure tone average (PTA) was calculated. The average PTA for the whole cohort was 16.4833 dB HL (SD = 7.3184, Figure 1a). Participants’ PTA values were positively correlated with their age (R^2^ = 0.2649, F(6.275, 114) = 42.44, p < 0.0001; Figure 1b). Participants also completed the Speech, Spatial, and Quality of Hearing Scale (SSQ12), a self-report questionnaire of sound perception abilities in noisy environments. The average score was 7.237/10 (SD = 1.377), where higher scores indicate better perceptual abilities.

**Figure 1.**
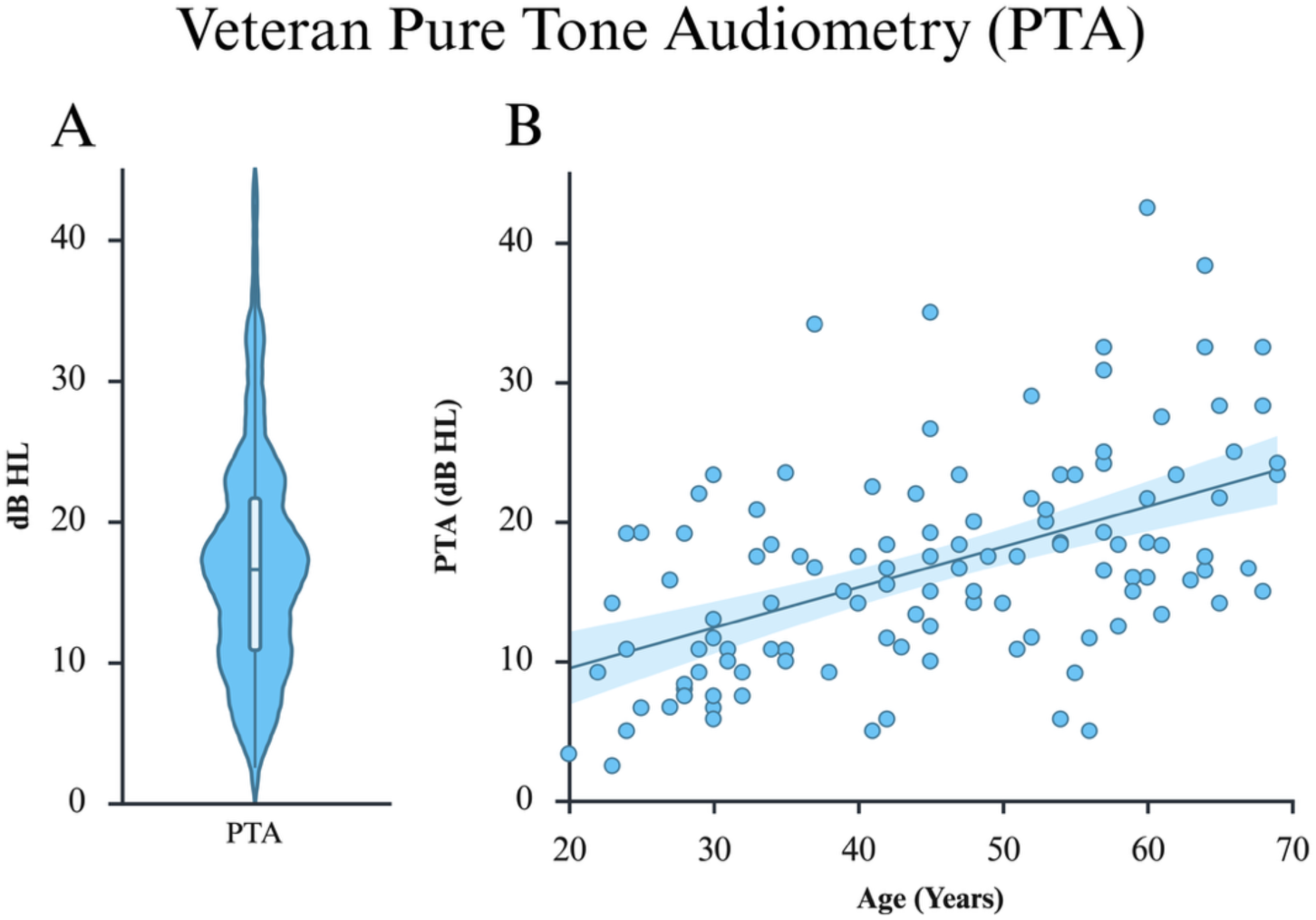

### Continuous Spatial Attention Task Performance

In our Cheech-based spatial attention task, accuracy color word hit was significantly higher for the speech-in-quiet (SIQ) conditions than the speech-in-noise (SIN) conditions as expected (t(113) = −5.511, p = <0.0001; SIQ M = 89.78 ± 8.3892% SD, SIN M = 81.977 ± 11.9243% SD, respectively; Figure 2a). The reaction time (RT) across conditions was also significantly faster for SIQ compared to the SIN condition (t(113) = −5.6789, p = <0.0001; SIQ M = 0.8577 ± 0.1198 seconds, SIN M = 0.9159 ± 0.1215 seconds; Figure 2b). The participants also scored significantly higher for the comprehension questions during the SIQ condition (M = 78.506, SD = 26.6238) when compared to the SIN condition (M = 66.7956, SD = 26.959, Figure 2c).

**Figure 2.**
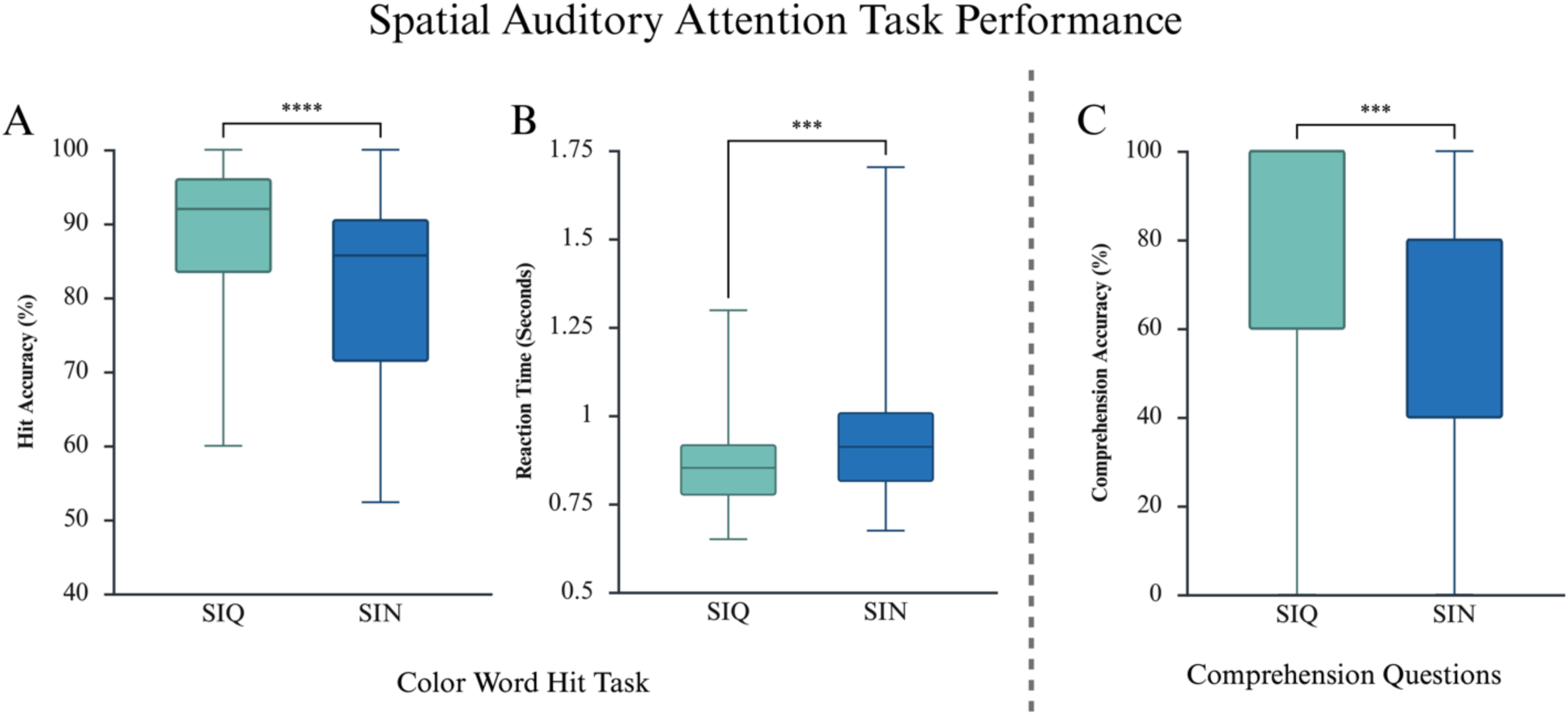

### Relationships between cognitive, audiologic, and behavioral speech listening measures

#### Reaction time

Mixed effects linear multiple regression was used to investigate the relation of hit RT and our predictor variables (age, PTA, SSQ, MoCA, and listening condition; Table 1). Significant main effects included the listening condition (SIQ vs SIN) and age as well as an interaction between participant age and condition. Figure 3a shows the predicted relationship between RT and age across both conditions. Marginal means demonstrated no significant trend during the SIQ conditions (β = −0.0127, SE = 0.0132, p = 0.3360), while a significant positive association between age and RT was noted for the SIN condition (β = 0.0275, SE = 0.0132, p = 0.0378).

**Figure 3.**
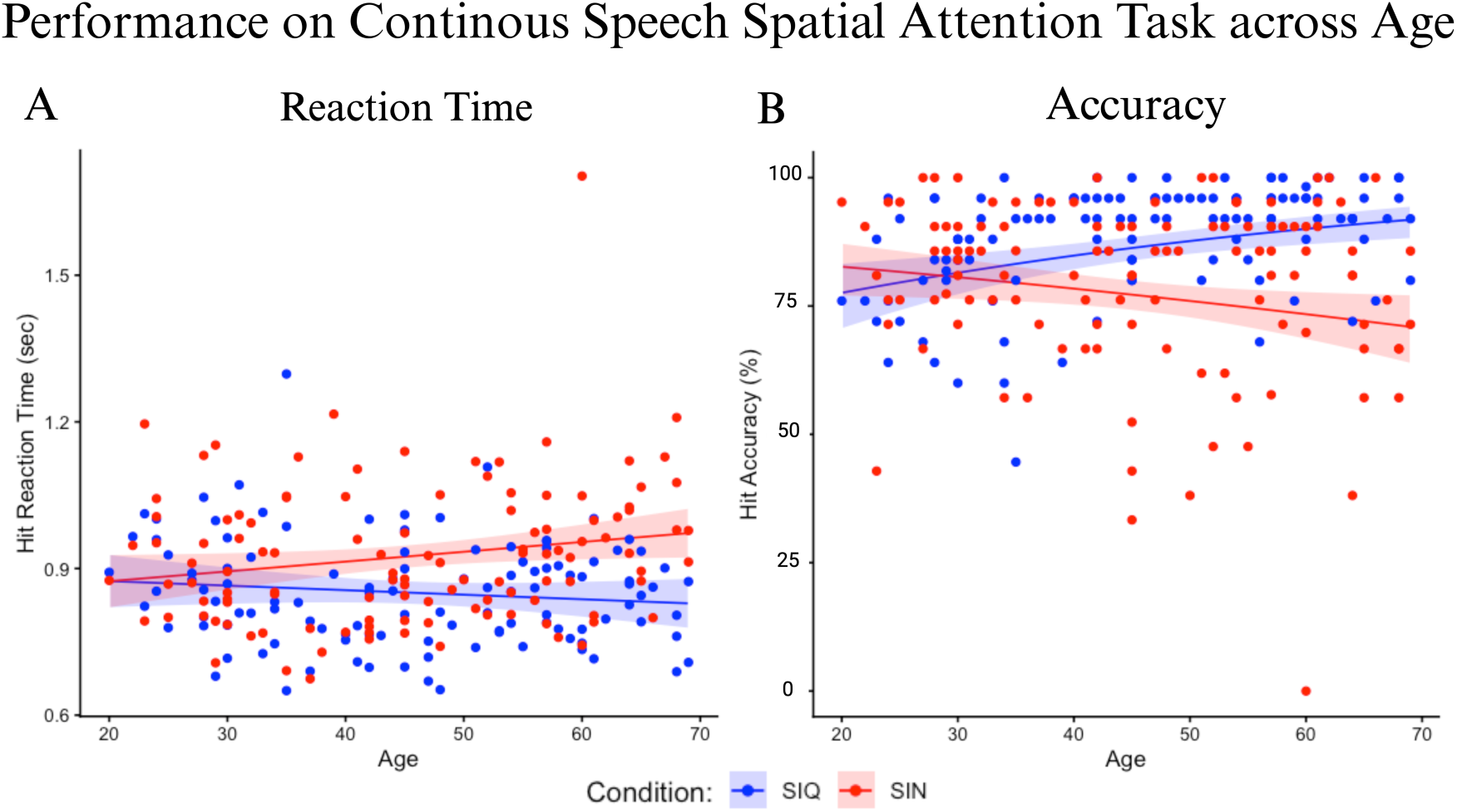

**Table 1.** Mixed Effect Linear Multiple Regression Model to Predict Hit Reaction Time.

| Predictor Variable | Estimate | SE | Z | p |
| --- | --- | --- | --- | --- |
| Intercept | 0.925214 | 0.011620 | 79.62 | **< .0001** |
| Condition | -0.074696 | 0.013108 | -5.70 | **< .0001** |
| Age | 0.027466 | 0.013220 | 2.08 | **0.03776** |
| PTA | -0.001135 | 0.011609 | -0.10 | 0.92214 |
| SSQ | -0.000886 | 0.008991 | -0.10 | 0.92151 |
| MoCA | 0.003813 | 0.010470 | 0.36 | 0.71573 |
| Condition:Age | -0.040184 | 0.013165 | -3.05 | **0.00227** |

That is, RT increases with the participants’ age when there is a competing stimulus, but this trend is not observed when listening to speech with no competing stimulus.

#### Identification accuracy

Mixed effects beta regression modeling was used to investigate the relationship between color word hit accuracy on the task and our predictor variables (Table 2). Significant main effects were observed for listening condition, age, and SSQ scores. In addition, two-way interactions between the following variables were found: condition & age, PTA & MoCA scores, and SSQ12 & MoCA scores. Three-way interactions were also observed between age, PTA, and MoCA as well as condition, SSQ, and MoCA scores.

**Table 2.** Mixed Effect Beta Multiple Regression Model to Predict Hit Accuracy.

| Predictor Variable | Estimate | SE | Z | p |
| --- | --- | --- | --- | --- |
| Intercept | 1.235047 | 0.082618 | 14.949 | < .0001 |
| Condition | 0.625851 | 0.099981 | 6.260 | < .0001 |
| Age | -0.182047 | 0.083984 | -2.168 | **0.03019** |
| PTA | 0.004266 | 0.076880 | 0.055 | 0.95575 |
| SSQ | 0.237271 | 0.074766 | 3.174 | **0.00151** |
| MoCA | 0.060863 | 0.085786 | 0.709 | 0.47803 |
| Condition:Age | 0.514731 | 0.099009 | 5.199 | < .0001 |
| Condition:SSQ | -0.175256 | 0.090729 | -1.932 | 0.05340 |
| Condition:MoCA | 0.096543 | 0.099656 | 0.969 | 0.33266 |
| Age:PTA | 0.021132 | 0.065672 | 0.322 | 0.74762 |
| Age:SSQ | 0.074824 | 0.055713 | 1.343 | 0.17926 |
| Age:MoCA | -0.169196 | 0.090071 | -1.878 | 0.06032 |
| PTA:MoCA | 0.201908 | 0.089294 | 2.261 | **0.02375** |
| SSQ:MoCA | 0.275985 | 0.068523 | 4.028 | < .0001 |
| Age:PTA:MoCA | -0.192701 | 0.088459 | -2.178 | **0.02937** |
| Condition:SSQ:MoCA | -0.190924 | 0.079474 | -2.402 | **0.01629** |

Figure 2b shows the relationship between the participants’ hit accuracy and their age across condition. It is seen that the SIQ conditions performance has a significant positive trend with the participant age (β = 0.328, SE = 0.0940, p = 0.0005), while there is a significant negative trend for the SIN conditions and age (β = −0.187, SE = 0.0839, p = 0.0260).

There is a main effect between SSQ12 scores and hit accuracy. When performance for each condition is isolated, there is no significant trend between SIQ accuracy and SSQ12 scores (Figure 4, β = 0.328, SE = 0.0940, p = 0.0005). SIN hit accuracy has a significant positive trend with SSQ12 scores (β = 0.2337, SE = 0.0745, p = 0.0017).

**Figure 4.**
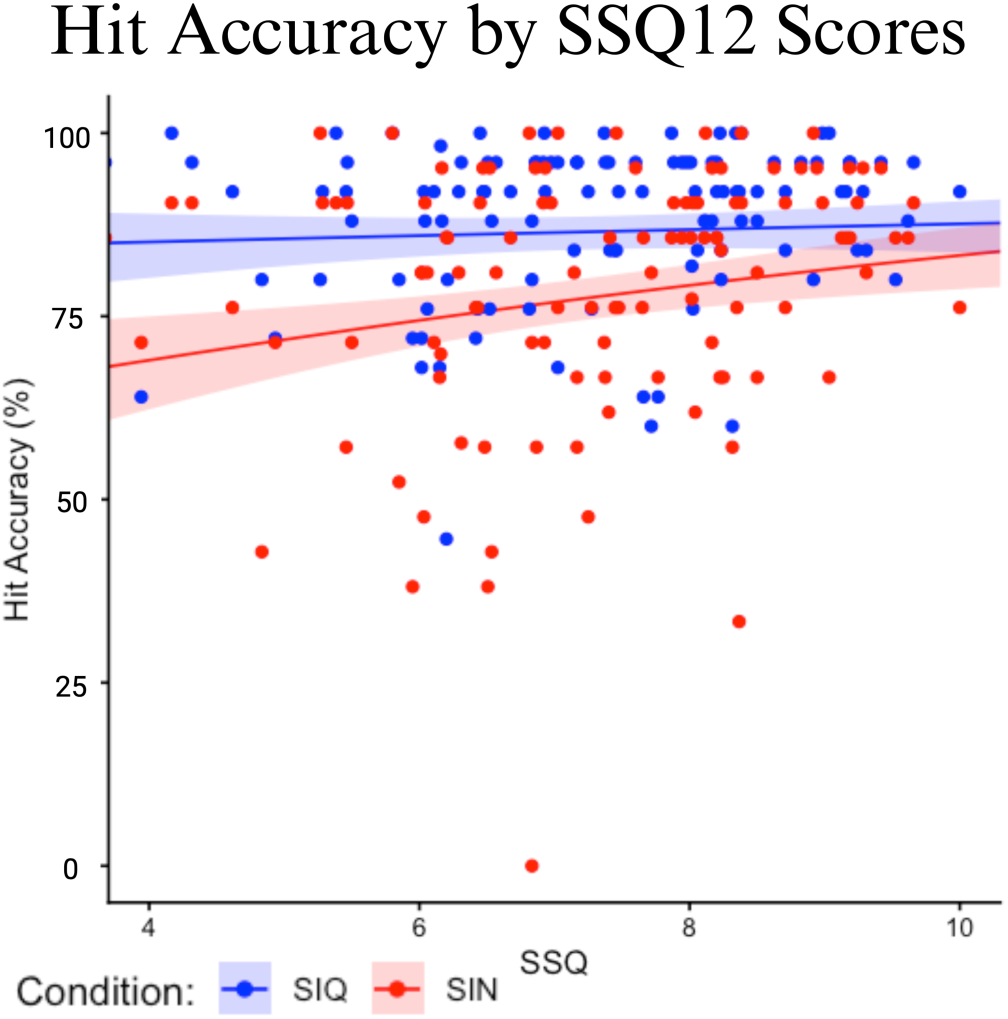

To break down the complex 3-way interaction between age, PTA, and MoCA scores, we conducted a series of spotlight analyses highlighting the underlying 2-way effects. Figure 5 represents the relationship between hit accuracy and PTA when a participant’s MoCA score is held at specific values (age held constant). The central plot (figure 5c) shows the relationship at the average MoCA score (M= 26.96) with each concentric plot for 1 standard deviation (figure 5b & figure 5d) and 2 standard deviations (figure 5a & figure 5e) away from the mean (SD = 1.614), respectively. Performance on the SIN condition is lower than performance on the SIQ conditions in each plot, showing a consistently better performance when completing the task with no competing, masking stimuli. According to our marginal means analysis, the predicted models for the lowest MoCA scores (∼24/30, Figure 5A) showed a significant negative trend where hit accuracy decreases with increasing PTA (SIQ & SIN trends: β = −0.3996, SE = 0.1910, p = 0.0367). Models for MoCA at 1 SD from the mean (∼25/30) show a nonsignificant but negative trend (SIQ & SIN trends: β = −0.1976, SE = 0.1150, p = 0.0863). Participants with average MoCA scores (∼27/30) show no significant association between PTA and hit accuracy (SIQ & SIN trends: β = 0.00427, SE = 0.0769, p = 0.9558). The predicted models where the MoCA scores are higher than the average score show a positive relationship between hit accuracy and PTA. At 1 SD from the mean (∼29/30), this positive trend is not significant (SIQ & SIN trends: β = 0.2062, SE = 0.1200, p = 0.0868) but it is significant at 2 SD away from the mean (∼30/30, SIQ & SIN trends: β = 0.4081, SE = 0.1980, p = 0.0388) This means that participants with worse hearing ability (high PTA) perform better than participants with better hearing ability if their rapid cognitive assessment is above average. Thus, hit accuracy performance decreases with worsening hearing ability (higher PTA) in those with lower-than-average cognitive ability whereas accuracy increases with perceptual thresholds in those with higher-than-average cognition, and this effect is stronger with more extreme MoCA scores (i.e., steeper slopes for 2 SD away from them mean than 1 SD away from the mean).

**Figure 5.**
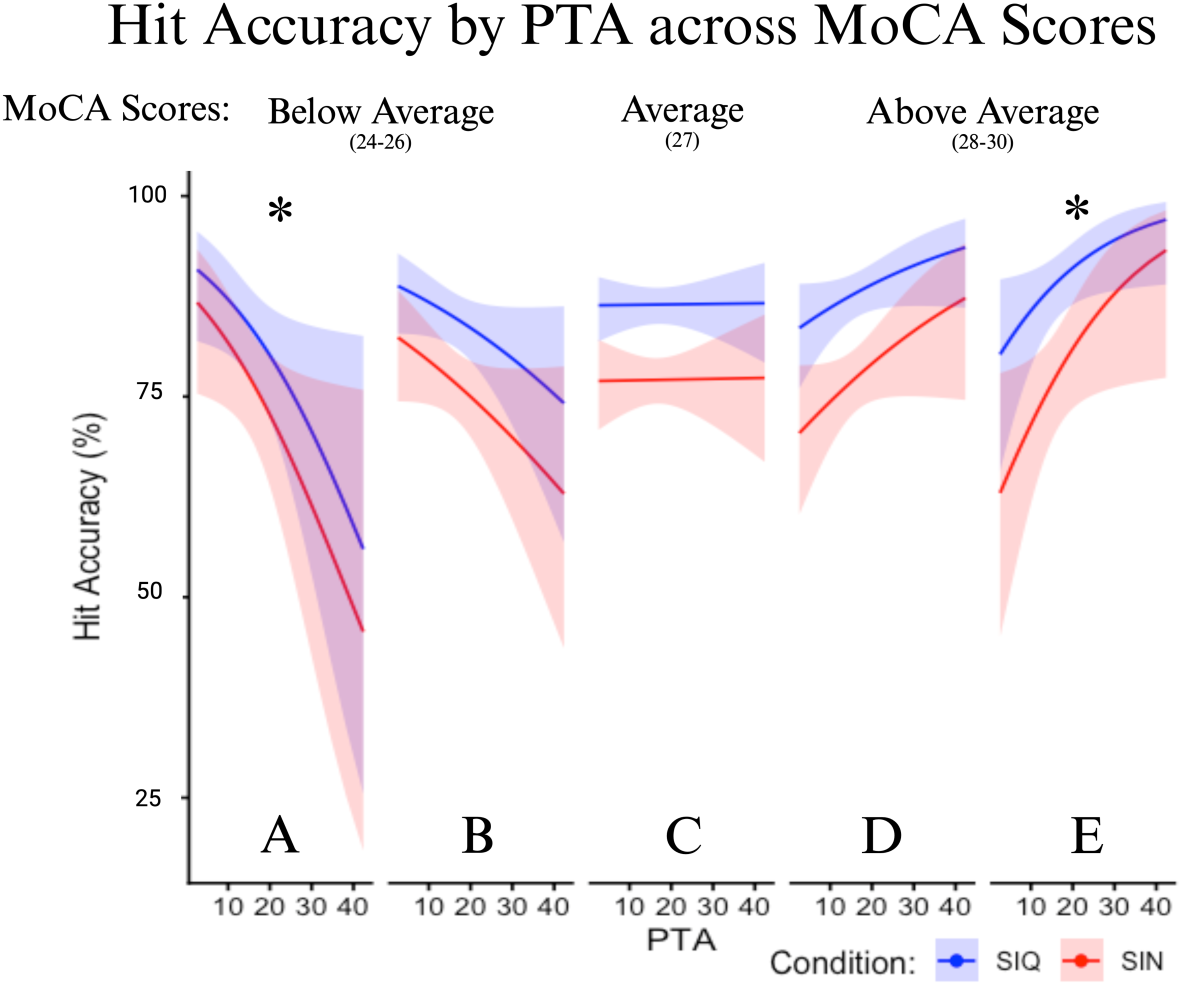

Figure 6 indicates the spotlight analysis results between hit accuracy, age, and MoCA scores (PTA held constant). In the SIQ condition, increasing age is generally associated with better hit rate accuracy, but this effect is only significant for those with average or below-average MoCA scores (−2 SD: β = 0.6652, SE = 0.2020, p = 0.0010; −1 SD: β = 0.4955, SE = 0.1290, p = 0.0001; mean: β = 0.3258, SE = 0.0940, p = 0.005; +1 SD: β = 0.1562, SE = 0.1310, p = 0.2345; +2 SD: β = −0.0135, SE = 0.2050, p = 0.9473). Meanwhile, increasing age is associated with worse accuracy in the SIN condition accuracy for average and above-average MoCA scores (−2 SD: β = 0.1505, SE = 0.1970, p = 0.4441, -1 SD: β = −0.0192, SE = 0.1210, p = 0.8742; mean: β = - 0.1889, SE = 0.08939, p = 0.0244; +1 SD: β = −0.3586, SE = 0.1250, p = 0.0041; +2 SD, β = - 0.5283, SE = 0.2010, p = 0.0086). Collectively, these results suggest that advancing age is associated with worse SIN listening abilities, particularly for those with average or above-average cognition, whereas age has a paradoxically *beneficial* role for target word identification in quiet among those with cognitive abilities in the average or below-average range.

**Figure 6.**
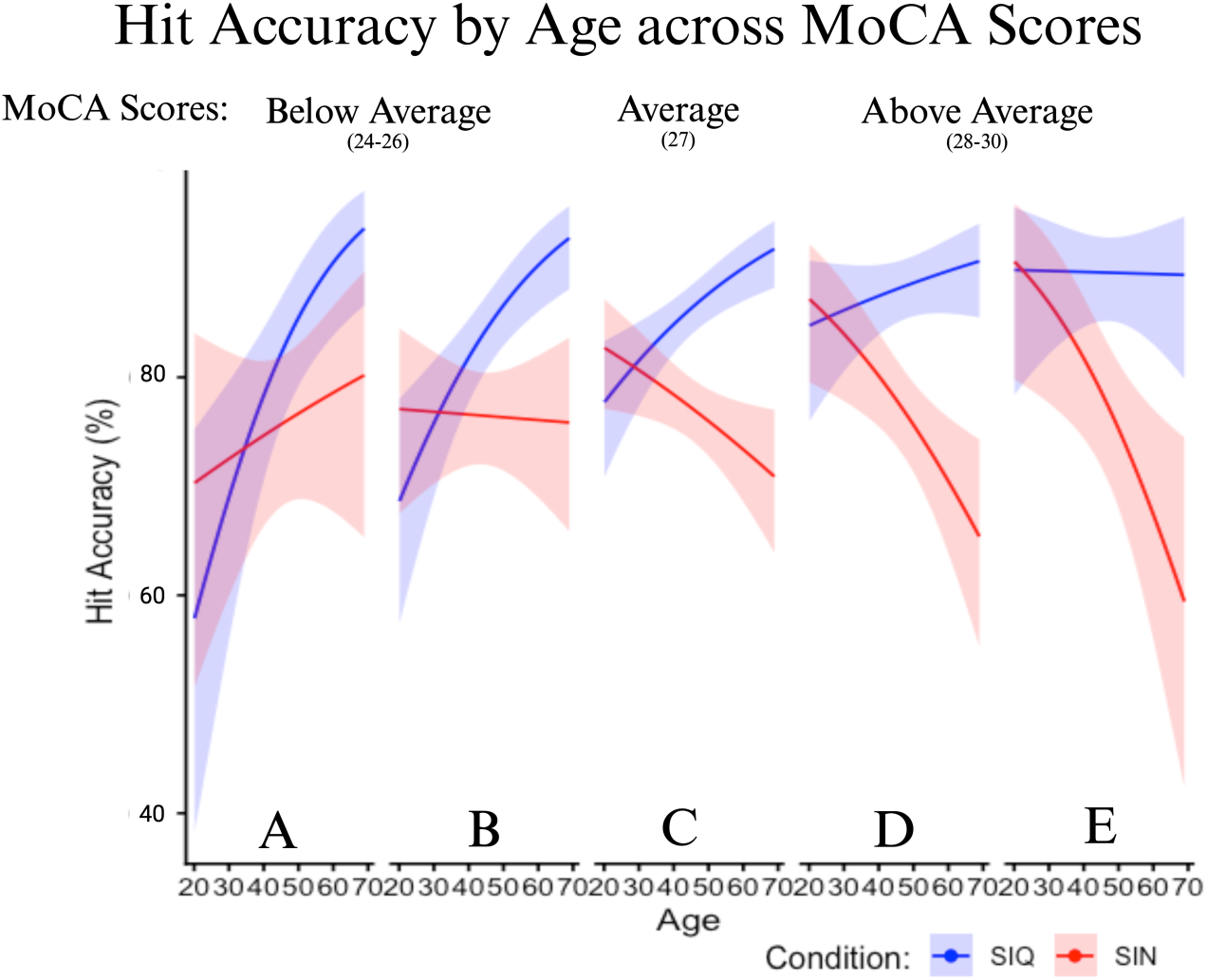

#### Narrative Comprehension Performance

After completing the spatial selective attention task, participants’ comprehension of the target narrative was evaluated with 5 multiple choice questions. The participants’ average accuracy for those questions was modeled using the same predictor variables (PTA, age, SSQ, and MoCA) in a mixed effects beta multiple regression model (Table 3). Model results suggested a main effect of listening condition and MoCA scores as well as an interaction between condition and PTA.

**Table 3.**
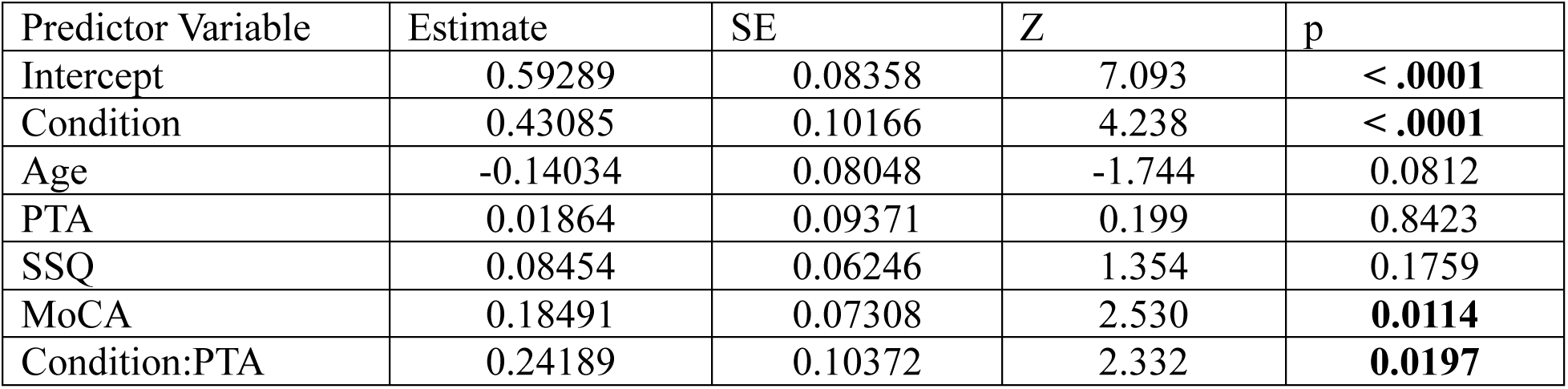
Mixed Effect Beta Multiple Regression Model to Predict Comprehension Accuracy.

Figure 7a shows the significant positive relationship between comprehension accuracy and MoCA ability for both conditions (overall β = 0.185, SE = 0.0731, p = 0.0114). Figure 7b plots participants’ comprehension performance across PTA values and condition. SIQ comprehension performance significantly increases with the participant’s age (β = 0.2605, SE = 0.1010, p = 0.0095), while there is no significant trend for the SIN condition (β = 0.0186, SE = 0.0937, p = 0.8423).

**Figure 7.**
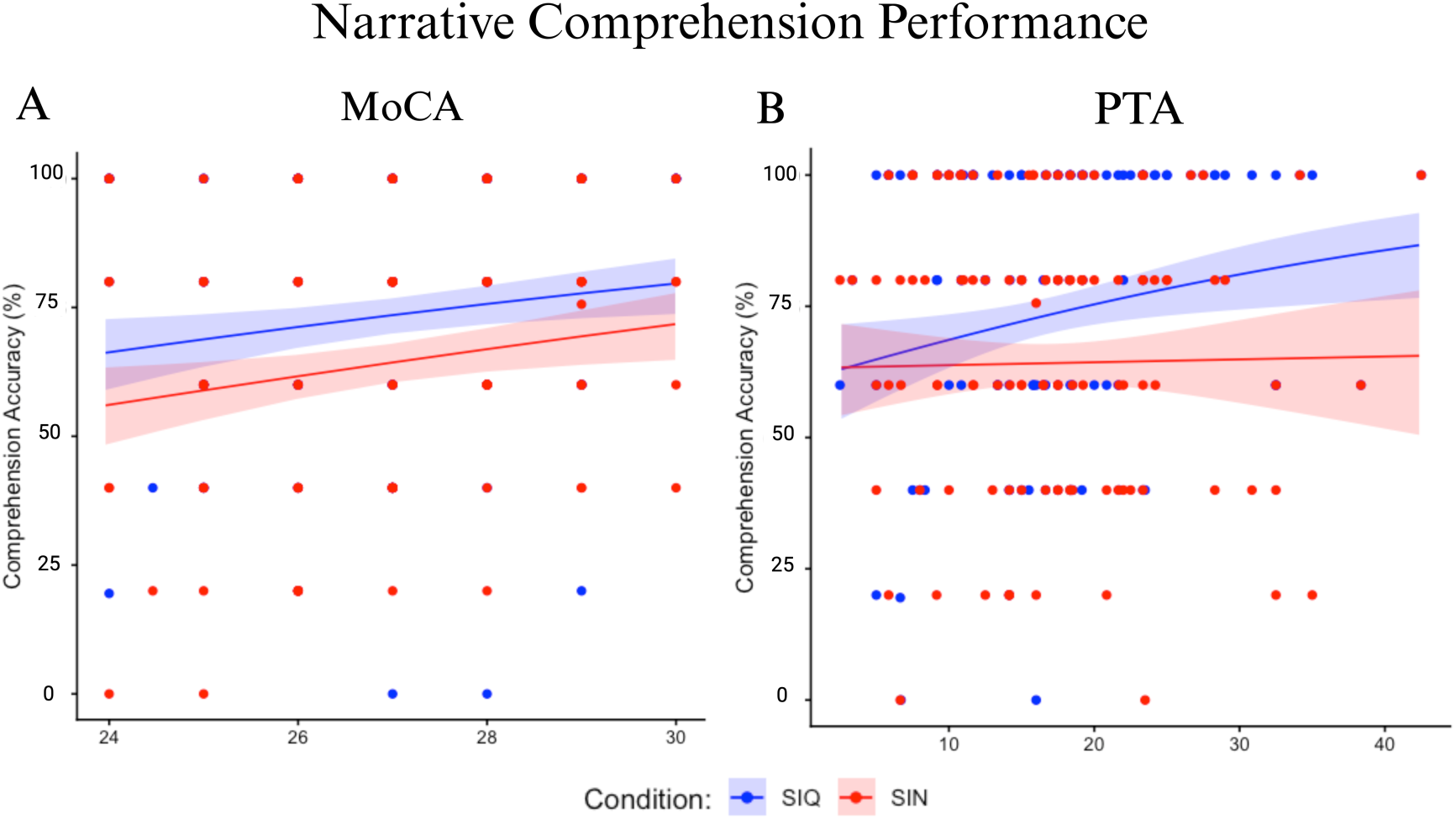

## Discussion

Our investigation evaluated whether pure tone audiometry (PTA) meaningfully predicts speech-in-quiet (SIQ) and speech-in-noise (SIN) perception in a cohort of veterans using an ecologically valid, spatial auditory attention task. We investigated two conditions: a mono-talker “quiet” condition with a single narrative stream and a dual-talker SIN condition containing a competing narrative masker designed to approximate everyday listening challenges. Performance was quantified via color word hit accuracy, color word reaction time (RT) and comprehension question accuracy. Complementary clinical and self-report measures including the MoCA and SSQ12 as well as participants’ age provided context for interpreting individual differences. Our findings demonstrate that that PTA, age, cognitive ability, and task condition interact in complex ways to shape auditory performance.

### Performance on the Novel Speech Attention Task

Participants performed significantly better and responded faster during target word identification in the SIQ condition compared to the SIN condition, confirming that the addition of a competitive talker introduces greater perceptual and cognitive demand (S. Anderson et al., 2013; Rönnberg et al., 2013). Greater variability in SIN accuracy and RT further reinforces the known challenge of processing speech in complex environments. Additionally, comprehension in the SIN was lower than in the SIQ condition, demonstrating that listening in noise reduces narrative encoding regardless of cognitive ability.

The effect of condition was not uniform across individuals. A significant condition x age interaction emerged for both accuracy and RT where younger participants performed comparably across SIQ and SIN conditions, while older participants showed a diverging trend of improved accuracy in the SIQ condition but reduced accuracy and slower RT in the SIN condition. This dissociation suggests that the cognitive and neural demands of selective attention under competing-speech conditions disproportionately affect listeners as they age. If age-related declines reflected only peripheral auditory changes, we would expect parallel degradation across both conditions. Instead, only the SIN condition was significantly impacted with increasing age while SIQ RT showed no significant age-related trend. Notably, SIQ accuracy actually improved with age, a pattern consistent with the idea that long-term language exposure may strengthen top-down linguistic representations and facilitate attention to a single narrative stream (Wingfield & Stine-Morrow, 2000). These benefits, however, appear to be lost once an additional, ecologically relevant competing speech stream is introduced. This mirrors subjective accounts from individuals with hidden hearing loss who frequently report no difficulty in quiet one-on-one settings but significant struggle in multi-talker environments (Spankovich et al., 2018; Tremblay et al., 2015). An important consideration here is that the additional cognitive and neural burden of attending to a target in noise stresses processing resources in ways that listening in quiet or more ideal conditions does not (Rönnberg et al., 2013).

### The Role of Age, Audiometric Sensitivity, and Cognitive Ability in Speech Perception

Collectively, our results highlight the role of cognitive abilities (as measured by the MoCA) for supporting speech listening abilities. For example, comprehension question accuracy generally increased with cognitive ability. This supports literature linking narrative and discourse comprehension with cognitive resources including working memory and executive function (S. Anderson et al., 2013; Rönnberg et al., 2013; Wingfield & Stine-Morrow, 2000). Importantly, this MoCA effect was not isolated to comprehension. The mixed-effects models revealed an informative set of interactions among age, MoCA scores, PTA, and condition.

Spotlight analyses stratified by MoCA score (mean = 26.96, SD = 1.6) revealed how cognitive ability modulates the relationship between audiometric and behavioral measures. Specifically, SIQ accuracy increased with age when participants exhibited below-average MoCA scores. For participants with average MoCA scores, SIQ accuracy increased with age while SIN accuracy declined. Meanwhile, SIQ accuracy showed no significant age-related trend while SIN accuracy declined most sharply when MoCA scores were high. One possible interpretation is that older adults with high cognitive ability may engage more effortful, top-down compensatory strategies in quiet conditions, benefiting from a lifetime of language exposure, that are overwhelmed once a competing speech stream is introduced. This aligns with frameworks suggesting that cognitive resources enable compensatory listening in straightforward environments but have limitations when put under complex conditions (Rönnberg et al., 2013; Wingfield & Stine-Morrow, 2000). However, the mechanism underlying steeper SIN decline in the highest MoCA group warrants further investigation. A significant age x PTA x MoCA interaction further demonstrated that the relationship between age, hearing thresholds, and behavioral performance is not straightforward and depends critically on cognitive ability as well. These interactions highlight that aging effects on speech perception are heterogeneous across individuals, driven not just by peripheral auditory sensitivity but by a more complex interplay of perceptual, cognitive, and even experiential factors.

A particularly interesting finding emerged from the PTA x MoCA interaction on hit accuracy. When stratified by MoCA score, the relationship between PTA and task performance reversed direction across cognitive ability groups. Participants with below-average MoCA scores showed an expected pattern of worse hit accuracy being associated with higher (poorer) PTA thresholds. This reflects general intuition about this where lower auditory sensitivity reduces the quality of the incoming signal leading to impaired speech recognition. However, participants with above-average MoCA scores showed the opposite where higher PTA was associated with better hit accuracy. This same trend appeared in comprehension data: SIQ comprehension accuracy increased as PTA worsened, while in the SIN condition, PTA did not significantly predict comprehension. Taken together, these patterns across both hit accuracy and comprehension suggest that listeners with strong cognitive ability may be engaging compensatory mechanisms (such as enhanced linguistic prediction, selective attention, or working memory) that not only offset peripheral hearing limitations but also appear to provide a performance benefit in the context of this naturalistic task. In noisy conditions, these compensatory resources appear insufficient to overcome the added demands of selective attention and competing speech, reducing PTA’s relevance as a predictor. A significant age x PTA x MoCA three-way interaction further confirmed that the relationship among age, hearing thresholds, and behavioral performance is not straightforward and depends critically on cognitive ability. These findings collectively reinforce the view that cognition is a critical, and often overlooked, contributor to both SIQ and SIN perception.

### Limited Predictive Scope of PTA in Real-World Listening

A central finding of this study is the dissociation between PTA’s ability to predict to performance under quiet versus noisy listening conditions. For example, comprehension accuracy increased as PTA worsened in the SIQ condition, but PTA did not significantly predict comprehension accuracy for the SIN condition. This dissociation is critical because in quiet settings, cognitive compensation may partially offset elevated hearing thresholds and even produce these counterintuitive relationships between audiometric and behavioral measures. In noisy conditions, however, these compensatory resources are insufficient to overcome the increased demands of selective attention and increasing speech streams, and PTA becomes a less relevant predictor of comprehension performance. Yet, the main effect of PTA was not significant for either hit accuracy or reaction time when condition, age, and cognitive ability were included in the model. The relationship between PTA and performance was instead dependent on cognitive ability (as captured by the PTA x MoCA and Age x PTA x MoCA interactions). Individuals with similar PTAs showed substantial variability in speech perception, reinforcing that PTA alone does not capture the complex interplay of auditory perception, cognitive resources, and attentional processes that truly shape real-world listening.

These findings align with a growing body of literature demonstrating that PTA, while clinically indispensable, provides limited insight into everyday listening challenges (Moore et al., 2014; Musiek et al., 2017; Pienkowski, 2017). PTA assesses audibility for isolated tones in quiet, a fundamentally low-demand task that does not incorporate temporal cues, informational masking, spatial attention, or higher-order processing. Many individuals with “normal” audiograms experience substantial SIN difficulties attributable to synaptopathy, cognitive factors, temporal coding deficits, or neural processing differences (Billings & Madsen, 2018; Coffey et al., 2017). This decoupling helps explain why PTA remains useful for identifying significant sensory loss (e.g., in older adults or candidates for hearing devices (Maeda et al., 2018)) but fails to account for functional impairments in listeners with mild or early-stage deficits. Because PTA captures only the auditory processes required to detect pure tones, it lacks the sensitivity to reflect complex, multifaceted perceptual demands of SIN, precisely the domains where listeners often report the greatest challenges.

### Validation With Perceived Speech Listening Difficulties

Another goal of this investigation was to determine whether our novel task paradigm aligns with the real-world experiences of individuals. Analyses demonstrated that our task captures perceptually meaningful variance in self-reported and behavioral hearing difficulty. To validate the SIN condition specifically, we examined relationships between task performance and SSQ12 self-report scores. We observed a significant negative relationship with SIN hit accuracy in which individuals who rated themselves as experiencing greater difficulty hearing in noisy everyday environments performed worse on our dual-talker SIN condition. The absence of this relationship in the SIQ condition particularly is informative in that it demonstrates that the SSQ12 covariance is specific to the noise condition rather than reflecting a general difference in speech processing ability or motivation.

### Implications for Clinical Practice and Assessment

The present findings underscore the need to expand audiological evaluation beyond pure tone thresholds, particularly for adults with normal to moderate hearing who report real-world listening difficulties. Functional measures such as SIN testing and brief cognitive screening can provide critical complementary information that PTA alone cannot. For middle-aged individuals especially, reliance solely on PTA risks underestimating functional communication challenges and mischaracterizing the underlying causes of listening difficulty. Our continuous-speech paradigm offers several advantages over traditional measures, including ecological validity, sensitivity to individual differences in both auditory and cognitive domains, and compatibility with neural recordings (EEG). Its naturalistic design may make it particularly well-suited for detecting hidden hearing loss, central auditory deficits, or cognitive contributors to listening difficulty, which are areas where standard clinical tools lack sensitivity. Integrating this task into multimodal, machine-learning (ML) diagnostic frameworks may ultimately improve identification of diverse hearing phenotypes and support more personalized, targeted intervention.

### Limitations and Future Directions

Several limitations should be noted. The veteran sample may differ from the general population in noise exposure history or auditory health, potentially limiting generalizability. Additionally, the SIN masker employed continuous competing speech rather than multitalker babble typically used in standardized assessments, increasing ecological validity but reducing direct comparability with existing tests. The unexpected relationships between PTA and performance, while interpretable as cognitive compensation, warrants replication and more detailed investigation, particularly regarding potential mediators such as listening strategy, cognitive reserve, or motivation. Future work should also continue integrating neural measures to elucidate why some listeners maintain strong SIN performance despite elevated audiometric thresholds or, conversely, why individuals with normal thresholds struggle in noise (Coffey et al., 2017; Liberman et al., 2016; Mankel et al., 2025; Shehabi et al., 2025).

## Conclusion

In summary, pure tone averages did not reliably predict speech-in-noise performance in either quiet or noise when age and cognitive ability were accounted for. Rather than operating as a simple linear predictor, PTA interacted with age and cognitive ability in complex ways, demonstrating that the relationship between audiometric sensitivity and real-world speech perception is fundamentally affected by cognitive factors. Together, these results highlight the limitations of relying solely on pure tone thresholds and support the incorporation of functional, ecologically grounded measures to more accurately assess communication ability and identify individuals at risk for hidden or central auditory deficits. The present task, sensitive to the combined influence of auditory, cognitive, and attentional processes, represents a valuable step toward a more comprehensive and clinically actionable characterization of real-world hearing ability.

## Acknowledgements

Funding for this research was provided by the Department of Defense (DoD) and the Child Family Fund for the Center for Mind & Brain. The authors thank the audiology and clinical research team of the University of California, Davis Health for conducting initial participant hearing evaluations including, Dr. Robert Ivory, Au.D., Dr. Mackenzie Quinn, Au.D., Dr. Rachel Krager, Au.D., Dr. Steven Zurawski, Au.D., Dr. Austin Childers, Au.D., Dr. Kimberly Smith, Au.D., Randev Sandhu, and Angela Beliveau. We also thank Cathleen Chan and Jillian McKie for their contributions to data collection, and Elyse Ehlert and Tiana Smith for their work in recruiting our participants. Additionally, we are grateful for valuable discussions with Sophie Burstein, Alicia Dye, Reina Itakura, Zacharay McNaughton, Ferdous Rahimi, Tyler Statema, Sana Shehabi, and Audrey Vargas. Most importantly, we extend our deepest gratitude to all of the participants whose time and effort made this research possible. This work is dedicated to the memory of our team member, lab mate, and friend, Karim Abou Najm.

## Author Contributions

D.C.C, K.M, H.B, D.S, and L.M.M designed research; N.E.W, B.B, and S.D performed research; D.C.C, K.M, B.B, N.E.W, R.W and L.M.M contributed analytic tools; B.B, N.E.W, and D.C.C analyzed data; N.E.W, D.C.C, K.M, B.B, R.W, S.D, and L.M.M wrote and edited the paper.

